# Plastome-based Phylogenomic analyses provide insights into the germplasm resource diversity of *Cibotium* in China

**DOI:** 10.1101/2023.03.21.533638

**Authors:** Ri-Hong Jiang, Si-Qi Liang, Fei Wu, Li-Ming Tang, Bo Qin, Ying-Ying Chen, Yao-Heng Huang, Kai-Xiang Li, Xian-Chun Zhang

**Affiliations:** Guangxi Key Laboratory of Special Non-wood Forest Cultivation and Utilization, Guangxi Engineering and Technology Research Center for Woody Spices, Guangxi Forestry Research Institute, Nanning, China; State Key Laboratory of Systematic and Evolutionary Botany, Institute of Botany, The Chinese Academy of Sciences, Beijing, China; China National Botanical Garden, Beijing, China; College of Life Sciences, University of Chinese Academy of Sciences, Beijing, China; Beijing Botanical Garden, Beijing, China; Beijing Floriculture Engineering Technology Research Centre, Beijing, China; Guangxi Forestry Industry Group Stock Corporation, Nanning, China

**Keywords:** chloroplast genome, *Cibotium barometz*, *Cibotium sino-burmaense*, conservation, DNA barcoding, endangered species, germplasm resource, species diversity

## Abstract

Germplasm resource is the source of herbal medicine production. Cultivation of superior germplasm resources helps to resolve the serious conflict between long-term population persistence and growing market demand by producing materials with high quality consistently. *Cibotium barometz* is the original plant of cibotii rhizoma (“Gouji”), a traditional Chinese medicine used in the therapy of pain, weakness, and numbness of lower extremity. Long-history use of *Cibotium* has rendered wild populations of this species declined seriously in China. Without sufficient understanding of species and lineage diversity of *Cibotium*, it is difficult to propose a targeted conservation scheme at present, let alone selecting high-quality germplasm resources. In order to fill such a knowledge gap, this study sampled *C. barometz* and relative species throughout their distribution in China, performed genome skimming to obtain plastome data, and conducted phylogenomic analyses. We constructed a well-supported plastome phylogeny of Chinese *Cibotium*, which showed that three species with significant genetic difference distributed in China, namely *C. barometz*, *C. cumingii*, and *C. sino-burmaense*, a cryptic species endemic to NW Yunnan and adjacent region of NE Myanmar. Moreover, our results revealed two differentiated lineages of *C. barometz* distributed in the east and west side of a classic phylogeographic boundary that probably shaped by monsoons and landforms in China. We also evaluated the resolution of nine traditional barcode loci, and designed five new DNA barcodes based on the plastome data which can discriminate all these species and lineages of Chinese *Cibotium* accurately. These novel findings integrated genetic basis will guide conservation planners and medicinal plant breeders to build systematic conservation plans and exploit germplasm resources of *Cibotium* in China.

## INTRODUCTION

Traditional Chinese medicine plays an indispensable role in the treatment of multiple diseases in China and other developing countries (Newman et al., 2008). Apart from the traditional usage, many medicinal plants, such as *Artemisia annua* L. (artemisinin, Tu, 2016), *Huperzia javanica* (Sw.) C. Y. Yang (Huperzine A, Zangara, 2003; it worthy to note that in many studies the plant was named as *Huperzia serrata* (Thunb.) Trevis., a native species only found in NE Asia which do not produce Huperziane A, Chen et al., 2021), and *Panax notoginseng* (Burk.) F. H. Chen (Notoginseng triterpenes, Huang et al., 2021), have also been found as the source of modern pharmaceuticals and generated increasing attention. Although China harbors abundant medicinal plant diversity, original species of many commonly used herbal medicines are facing the risk of population decline and even extinction under a growing demand (Chen et al., 2016). Germplasm resource is the core of medicine production (Ma and Xiao, 1998; Huang et al., 2008; Zhang and Jiang, 2021; Meng et al., 2023). Cultivation of specific high-quality germplasm resources will not only resolve present conflict between conservation and exploitation, but also ensure a steady production of high-quality medicine (Ma and Xiao, 1998; Chen et al., 2016). Therefore, clarifying genetic background and diversity is the basic and crucial step of achieving sustainable utilization of medicinal plants, and also provides implications for the collection, identification, evaluation and conservation of germplasm resources (Schoen and Brown, 1993; Ma and Xiao, 1998; Yu et al., 2013; Khoury et al., 2022).

*Cibotium barometz* (L.) J. Sm. is the original species of traditional medicine cibotii rhizoma (“Gouji” in Chinese), the processed rhizome of which can be used in the therapy of pain, weakness, and numbness of lower extremity (Chinese Pharmacopoeia Commission, 2020; Figure 1A). Phytochemical researches have showed that the extract of its rhizomes is rich in active compounds such as pterosins, terpenes, steroids, flavonoids, glucosides, phenolic acids, and pyrones (Xu et al., 2012). Bioactivity experiments support its effect including the treatment of osteoporosis and osteoarthritis, antioxidant and antimicrobial activities, as well as abirritation (Ju et al., 2005; Cuong et al., 2009; Zhao et al., 2011; Li et al., 2014; Fu et al, 2017; Heng et al, 2020; Sun, 2021). Pot cultures and crafts of this species are also popular on the market because of its elegant evergreen large fronds and stump-like rhizomes covered with long, soft, golden hairs resembling gold-hair dogs (Figure 1B- D). Medicinal and ornamental values have resulted in destructive plunder of abundant natural resources of *C. barometz* in China. Investigation has shown that uncontrolled collection and habitat deconstruction are major threats of its population survival (Zhang et al., 2002).

**FIGURE 1.**
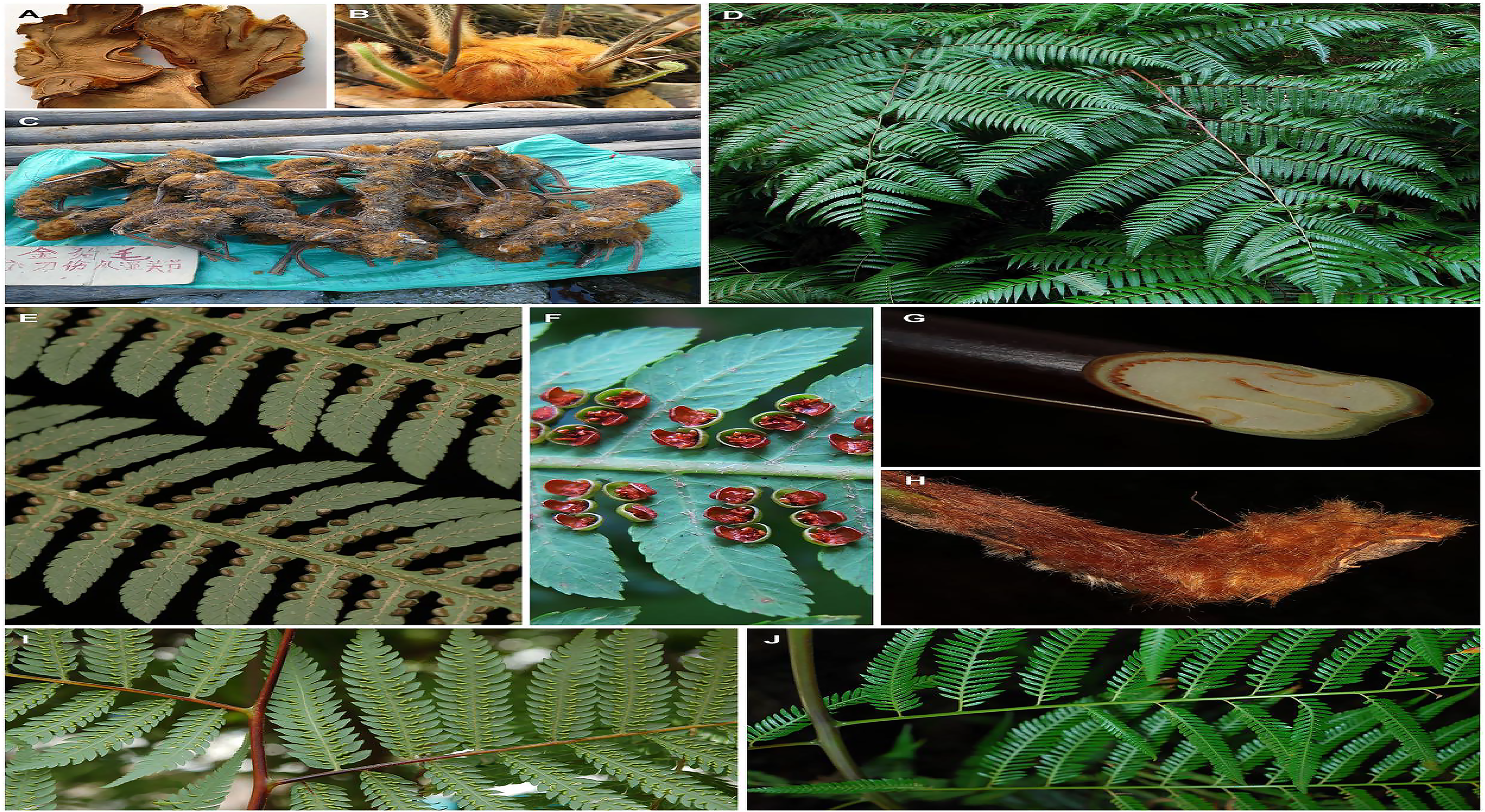
Morphology of *Cibotium* plants from China. **(A)** Dry sliced rhizomes of *C. barometz*, “Gouji”. (**B**) Rhizome, stipes and young fronds of *C. barometz* covered with golden filiform hairs. (**C**) Rhizomes of *C. barometz* sold as medicinal herbs at a village fair. (**D**) Fronds of *C. barometz*. (**E**) Veins, hairs and unopened sori on abaxial surface of pinnae of *C. borametz*. (**F**) Opened sori of *C. borametz*. (**G**-**H**) Cross section and basal part of stipe of *C. barometz*. **(I-J)** Basal part of pinna in *C. barometz* and *C. cumingii* showing the difference of basal pinnules on the basiscopic side. Photographs by R.-H. Jiang (A–B & D & F), X.-C. Zhang (C & I–J), and Q.-K. Ding (E & G–H).

*Cibotium barometz* is on the appendix II of CITES (Zhang et al., 2002; https://cites.org/eng/app/appendices.php). In the List of National Key Protected Wild Plants of China (State Forestry and Grassland Administration and the Ministry of Agriculture and Rural Affairs, P. R. China, 2021), the whole genus (with only two species known to China) is listed in Grade II Category. Although Chinese government has attached great importance to this genus, researchers are incapable of specifying which species or populations are key units awaiting conservation grounded in present knowledge. Such a phenomenon could lead to the waste of protective efforts and affect the maximization of medical value. Previous studies have showed that the genus *Cibotium* (Cibotiaceae, a member of the tree fern clade) comprises ca. 9-12 species distributed in tropical and subtropical regions of Asia, Central America and the Hawaiian Islands (Holttum, 1963; Palmer, 1994; Korall et al., 2006; Smith et al., 2006; Geiger et al., 2013), three Asian members of which form a monophyletic clade (Geiger et al., 2013). Two species, *A. barometz* and *A. cumingii* Kunze, are recognized from China (Zhang and Nishida, 2013), the former is widespread in southern China, northeastern India and extends to Malaysia, while the latter is only known from the Philippines, Ryukyu Islands, as well as Taiwan island of China (Holttum, 1963; Zhang and Nishida, 2013). However, the geographical pattern of genetic diversity and differentiation of *C. barometz* has not been explored throughout its wide distribution, let alone talking about the variation of medicinal values among different regions accurately.

In previous studies, several chloroplast DNA (cpDNA) fragments have been applied to phylogeny of the tree fern clade including *Cibotium* (Korall et al., 2006; Geiger et al., 2013). However, informative variation sites provided by these loci are too insufficient to illuminate the relationship within Chinese *Cibotium*. With the rapid development of next-generation sequencing (NGS) technologies, as well as advantages including low requirement of material quality, low costs and rich variable sites, chloroplast genome (plastome) has been used for phylogenetic reconstruction at different levels as well as species delimitation of closely related species in different plant lineages (e.g., Hammer et al., 2019; Wei and Zhang, 2020; Ji et al., 2021; Du et al., 2022; Xi et al., 2022; Zhang et al., 2022; Yang et al., 2023). Furthermore, plastomes can not only be applied to develop traditional DNA barcode but also used as a single genetic marker namely “ultra-barcode” (Nock et al., 2011; Kress et al., 2015; Hollingsworth et al., 2016), which largely benefits the identification of species or tissues lack of phenotypic divergence including products of medical plants (Park et al., 2021; Qin et al., 2022; Wang et al., 2022; Wei et al., 2022). In this study, we performed genome skimming and assembled the complete plastome of representative samples of *C. barometz* and relatives throughout the distribution range in China and adjacent areas. We aimed to 1) compare structure and composition variation on plastome among Chinese *Cibotium* species; 2) propose a phylogeny-based species delimitation; 3) investigate the geographical pattern of variation and diversity based on plastome data; and 4) suggest candidate barcodes for specific species and lineage identification of Chinese *Cibotium*.

## MATERIALS AND METHODS

### Taxon sampling, DNA extraction and Illumina sequencing

Frond tissues of 25 *Cibotium* individuals were collected for genome skimming throughout the distribution range of China and adjacent regions (Table 1, Figure 2). Most accessions were fresh fronds dried with silica-gel and preserved at 4℃ except five samples obtained from specimens deposited in the herbarium PE (Table 1). Based on the presence or lack of basal pinnules on basiscopic side of pinnae on voucher specimens (Figure 1I-J, Zhang and Nishida, 2013), samples were sorted into *C. barometz* or *C. cumingii* preliminarily.

**FIGURE 2.**
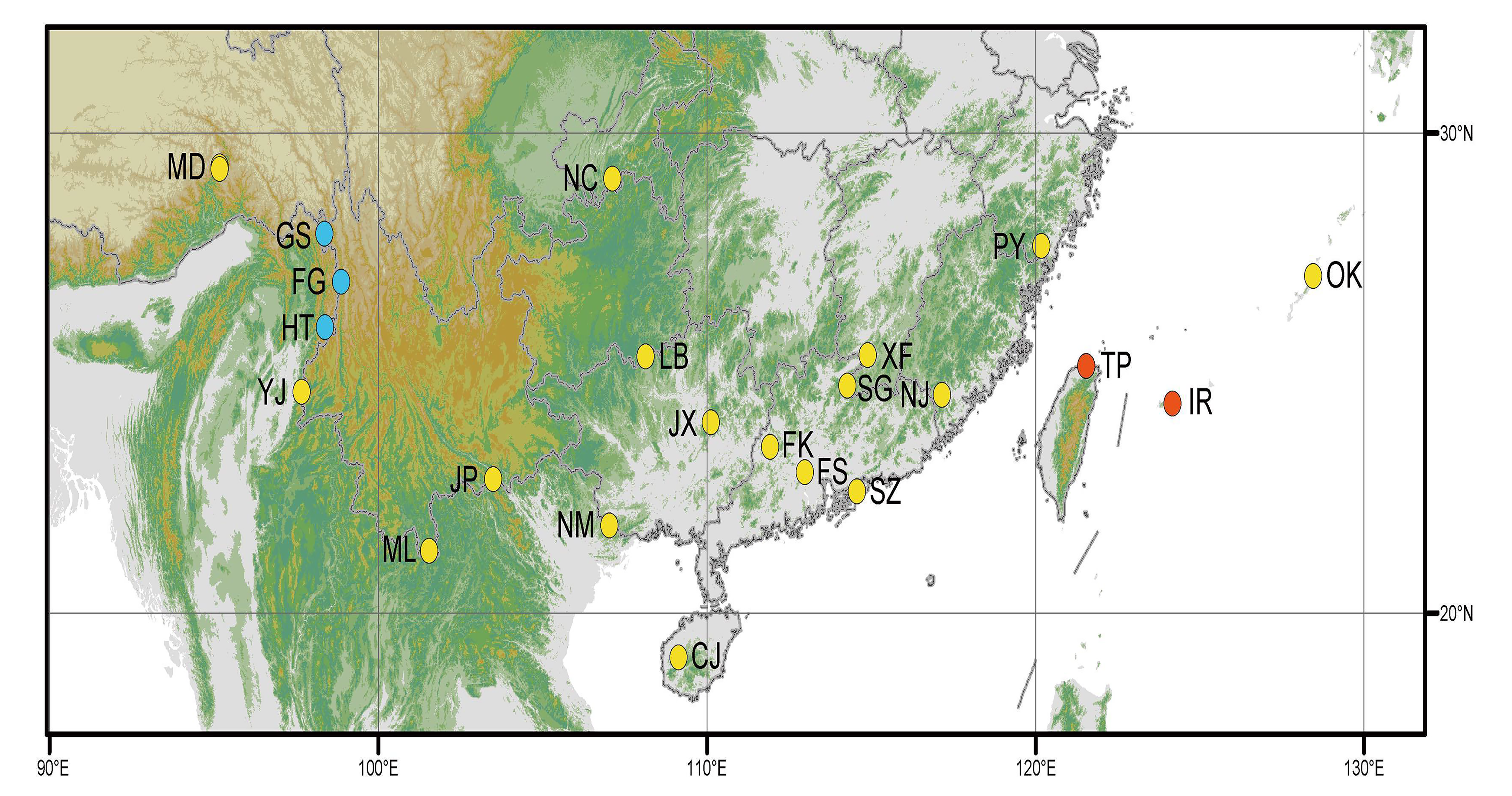
Map showing the distribution of *Cibotium* samples in this study. Yellow, blue and red dots represent localities of *C. barometz*, *C. sino-burmaense*, and *C. cumingii*, respectively. Code of each sampling locality means the same as Table 1.

**TABLE 1.**
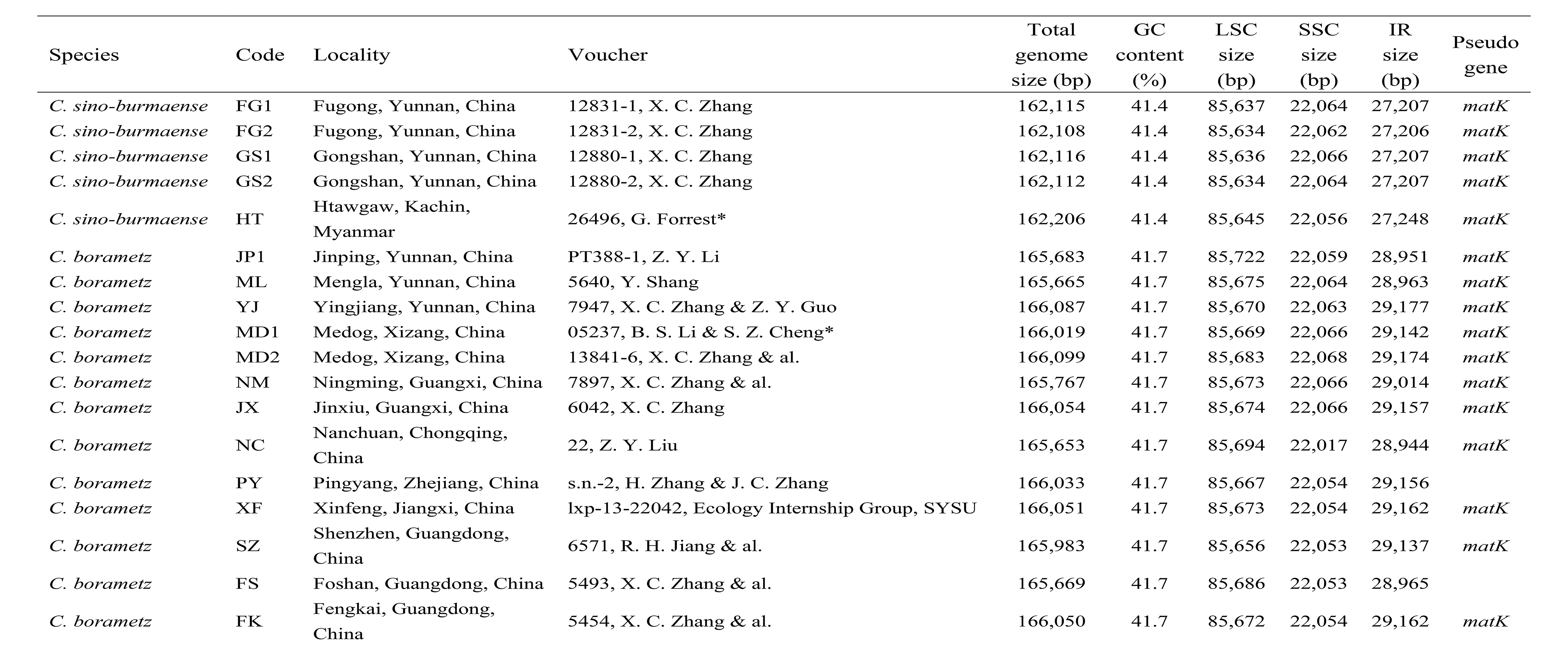
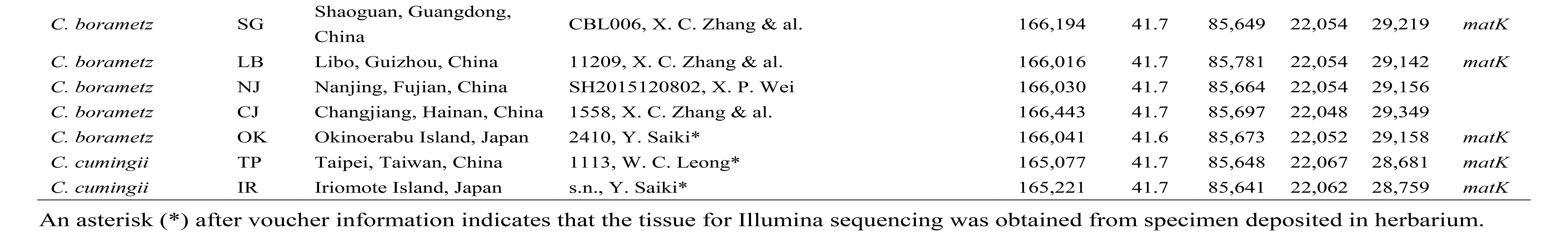
Summary of sampling information and plastome characteristics in this study.

All the tissue samples were sequenced at the Novogene Corporation (Beijing, China). Total genomic DNA was extracted with a modified CTAB procedure (Doyle and Doyle, 1987). Libraries with an insert size of 350 bp were constructed using a TruSeq Nano DNA HT Sample Preparation Kit (Illumina, San Diego, California, U.S.A.) following manufacturer’s recommendations. Paired-end reads (PE150) were then sequenced on an Illumina NovaSeq 6000 platform. After quality control of raw reads using ng_QC v.2.0 developed by Novogene Corporation with the default settings, we obtained ca. 2 to 4 Gb clean reads for each sample.

### Assembly and Annotation

We *de novo* assembled plastomes of all our samples with clean reads using the GetOrganelle toolkit (Jin et al., 2020) with recommended parameters. The complete plastome of *C. barometz* (NC_037893, Liu et al., 2018) downloaded from GenBank database was used as a reference during assembly and annotation. Assembly errors were identified in the initial assembly contigs and manually corrected by the mapping of raw reads to assembled sequences with Geneious v.11.1.4 (Kearse et al., 2012). Boundaries of large single-copy (LSC), small single-copy (SSC) and two inverse repeat regions (IRs) were detected using RepeatFinder v.1.0.1 (Volfovsky et al., 2001). Genome annotation was performed with GeSeq (Tillich et al., 2017) and Geneious v.11.1.4 (Kearse et al., 2012). Protein-coding sequences were checked against the National Center for Biotechnology Information (NCBI) database and manually corrected. tRNAs were confirmed with tRNAscan-SE v2.0.3 (Lowe and Chan, 2016). Final circular map of plastome were visualized with OGDraw v.1.3.1 (Greiner et al., 2019). We also used program LAGAN (Brudno et al., 2003) in mVISTA to compare the gene order and structure among different species with the plastome sequence alignment generated by MAFFT v.7.313 (Katoh and Standley, 2013).

### Phylogenetic analyses

The whole length plastome sequences of all *Cibotium* samples and the reference (NC_037893) as well as three outgroup species, i.e. *Alsophila spinulosa* (NC_012818), *Sphaeropteris brunoniana* (NC_051561) and *Plagiogyris euphlebia* (NC_046784), were aligned with MAFFT v.7.313 (Katoh and Standley, 2013) after the removal of one IR region. The alignment was then filtered using GBLOCKS v. 0.91b (Castresana, 2000) to remove ambiguously aligned regions. We also extracted protein-coding genes of each plastome with a python script (https://github.com/Kinggerm/PersonalUtilities/blob/master/get_annotated_regions_fr om_gb.py) and concatenated all these single gene alignments to build a protein-coding gene dataset for phylogenetic analyses. The best-fitting nucleotide substitution model of each alignment was determined based on Bayesian information criterion (BIC) by ModelFinder (Kalyaanamoorthy et al., 2017). Maximum likelihood (ML) analysis was performed with both datasets using IQ-TREE v.1.6.8 (Nguyen et al., 2015), with 10,000 ultrafast bootstrap replicates (Minh et al., 2013). Bayesian inference (BI) analysis was performed with the protein-coding gene dataset using MrBayes v.3.2.6 (Ronquist et al., 2012). One cold and three hot chains were run for 2,000,000 generations with sampling taken every 1,000 generations and a burn-in of 25%. The convergence of Markov chain Monte Carlo runs was checked with Tracer v.1.7.1 (Rambaut et al., 2018) to ensure that the effective sampling size (ESS) of all parameters were above 200. Phylogenetic trees were all visualized, rooted with *P. euphlebia* and edited in FigTree v.1.4.2 (Rambaut, 2014).

### Genetic diversity and divergence evaluation

With the whole length plastome sequence alignment including all 26 ingroup samples, we evaluated genetic diversity of each species and lineage (east and west lineage of *C. barometz*) by calculating nucleotide diversity (π) using DnaSP v.6.12.03 (Rozas et al., 2017). We also analyzed the number of fixed site differences and the average number of nucleotide substitutions per site (Dxy) between pairwise-species or lineages to show their divergence level.

### Candidate Barcoding Regions Detection and Verification

To identify candidate regions for species and even lineage discrimination in Chinese *Cibotium* plants, we first used DnaSP v.6.12.03 (Rozas et al., 2017) to evaluate π of the plastome sequence alignment of *C. barometz* with a window length of 800 bp and a step size of 200 bp. Nucleotide polymorphism sites fixed in specific species and lineage were also identified by checking the alignment including all *Cibotium* samples. Additionally, the feasibility and convenience of PCR amplification in practice was also taken into consideration, therefore, the chosen barcode regions are all shorter than 800 bp in length and have conservative flanks suitable for primers to combine with. Candidate loci meeting all these requirements were finally selected, PCR primers of which were designed with Primer3 v.2.3.7 (Koressaar & Remm, 2007; Untergasser & al., 2012).

We extracted sequences of newly selected loci and nine cpDNA markers (*atpA*, *atpB*, *rbcL*, *rps4*, *rbcL-accD*, *rbcL-atpB*, *trnG-trnR*, *trnL-trnF* and *rps4-trnS*) applied in previous studies (Korall et al., 2006; Geiger et al., 2013) from all our samples, included other accessible data of *C. barometz* and *C. cumingii* on GenBank, and aligned them. We counted the number of variable sites with MEGA v.10.1.6 (Kumar et al., 2018) and performed ML analysis on each alignment including outgroups following the same procedure as mentioned above. We compared topologies of resulted phylogenetic trees to the one built with plastome dataset to evaluate the efficiency of these loci in species and lineage discrimination. Multiple individuals of a specific taxon resolved as monophyletic with bootstrap support over 50% were treated as successfully discriminated.

## RESULTS

### Plastome Characteristics of *Cibotium*

Complete chloroplast genomes of 25 sampled individuals of *Cibotium* were obtained and assembled into circular molecules compromising one LSC, one SSC and two IRs (Figure 3, Table 1), which are all typical quadripartite structures. Complete plastomes of *C. cumingii* and the majority of *C. barometz* ranged from 165,077 to 166,443 bp in length with very similar GC contents ca. 41.7%, except five “*C. barometz*” samples probably of an unknown species collected from NW Yunnan and NE Myanmar with significantly shorter length (162,108–162,206 bp) and lower GC content (41.4 %). The length of LSC (85,634–85,781 bp) and SSC (22,017–22,067 bp) are rather stable among all accessions, whereas IR size varied among samples of *C. cumingii* (28,681– 28,759 bp), most *C. barometz* (28,944–29,349 bp) and those Yunnan-Myanmar samples (27,206–27,248 bp) with clearly discrete distribution. The boundaries of IR are exactly the same among all samples without any expansion or contraction. In comparison, the intergeneric regions between *rrn16* and *rps12* varied seriously among species (Figure 4), which mainly contributed to the IR size variation.

**FIGURE 3.**
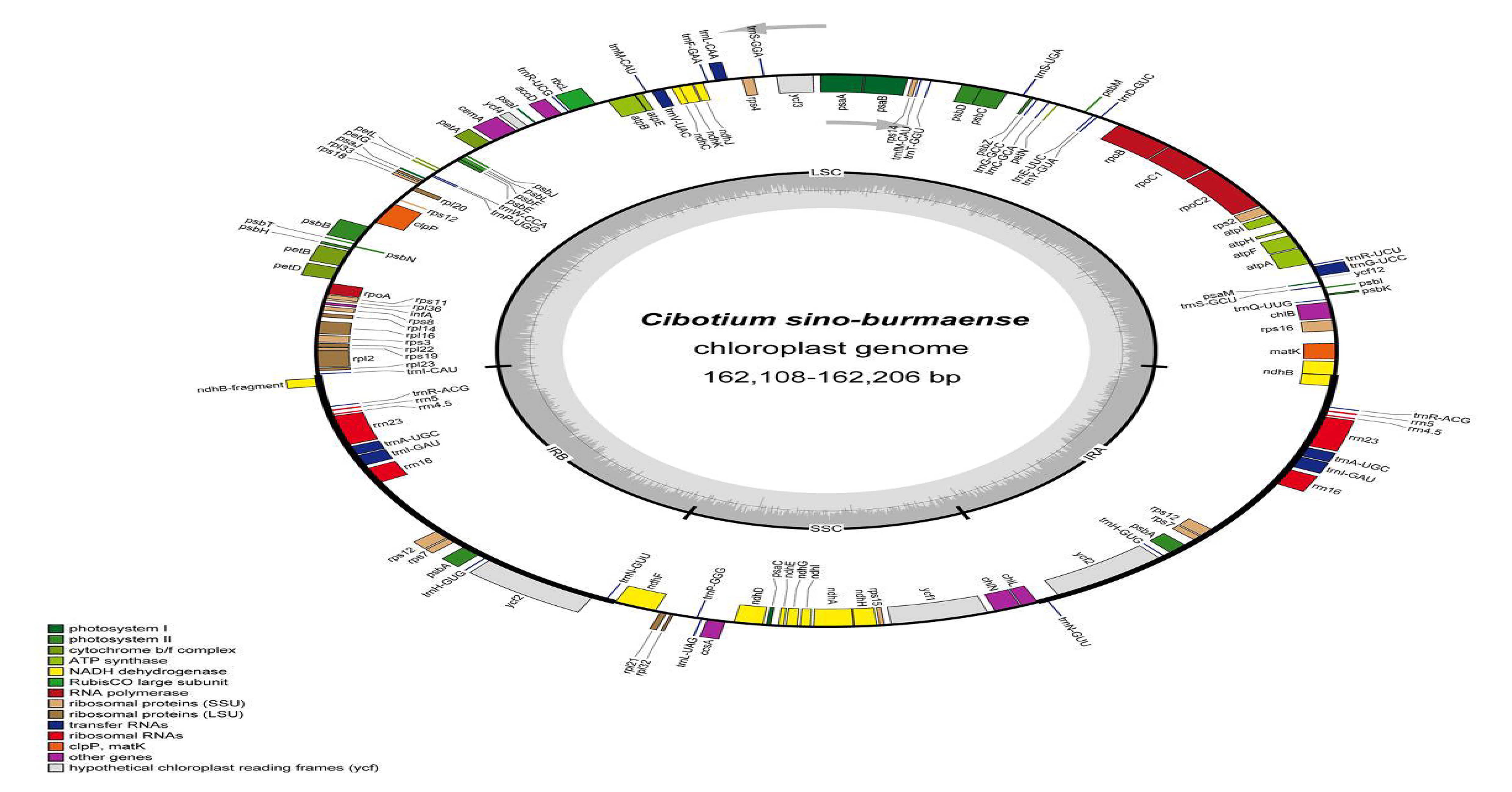
Plastome map of *Cibotium sino-burmaense*. Arrows indicate the direction of gene transcription. The dark grey area of the inner circle shows GC content variation among different region of the plastome.

**FIGURE 4.**
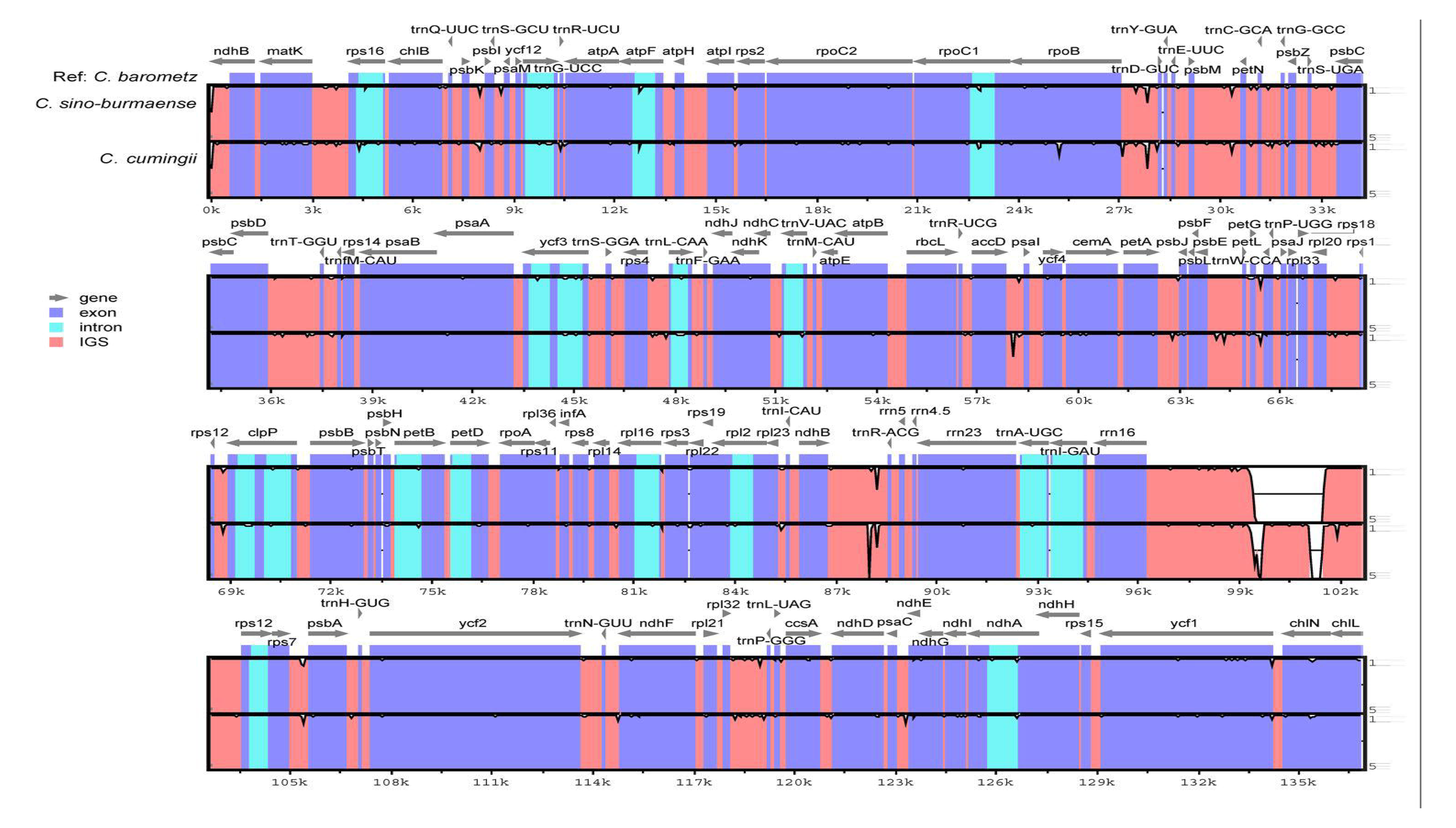
Sequence identity plot comparing plastome sequence and constitution of three Chinese *Cibotium* species with *C. barometz* as a reference. Each sequence starts with the beginning of LSC and end at the end of SSC. Gray arrows indicate genes with their orientation. A cut-off of 50% identity was used for the plot, and the Y-axis represents the percent identity ranging from 50% to100%.

All the plastomes encoded a total of 117 unique genes in identical order, including 85 protein-coding genes, 28 tRNA genes, four rRNA genes (Table 2, Figure 3), which is generally in consistent with the reference (Liu et al., 2018). In most samples of *C. barometz*, the annotated *matK* gene region could not be translated into protein successfully (pseudogenization) because of an early termination resulted from the missing of 1 or 2 nucleotides (Table 1). The gene *ycf2*, which was predicted as pseudogene (6,250 bp) due to a code shift mutation in the reference plastome of *C. barometz* (NC_037893, Liu et al., 2018), is found to be normal (6,249 bp) in all the samples of this study. All four rRNA genes, five tRNA genes (t*rnA-UGC, trnH-GUG*, *trnI-GAU*, *trnN-GUU*, *trnR-ACG*), and three protein coding genes (*rps7*, *psbA*, *ycf2*) are totally duplicated, whereas *ndhB* and *rps12* have one incomplete duplication merely.

**TABLE 2.**
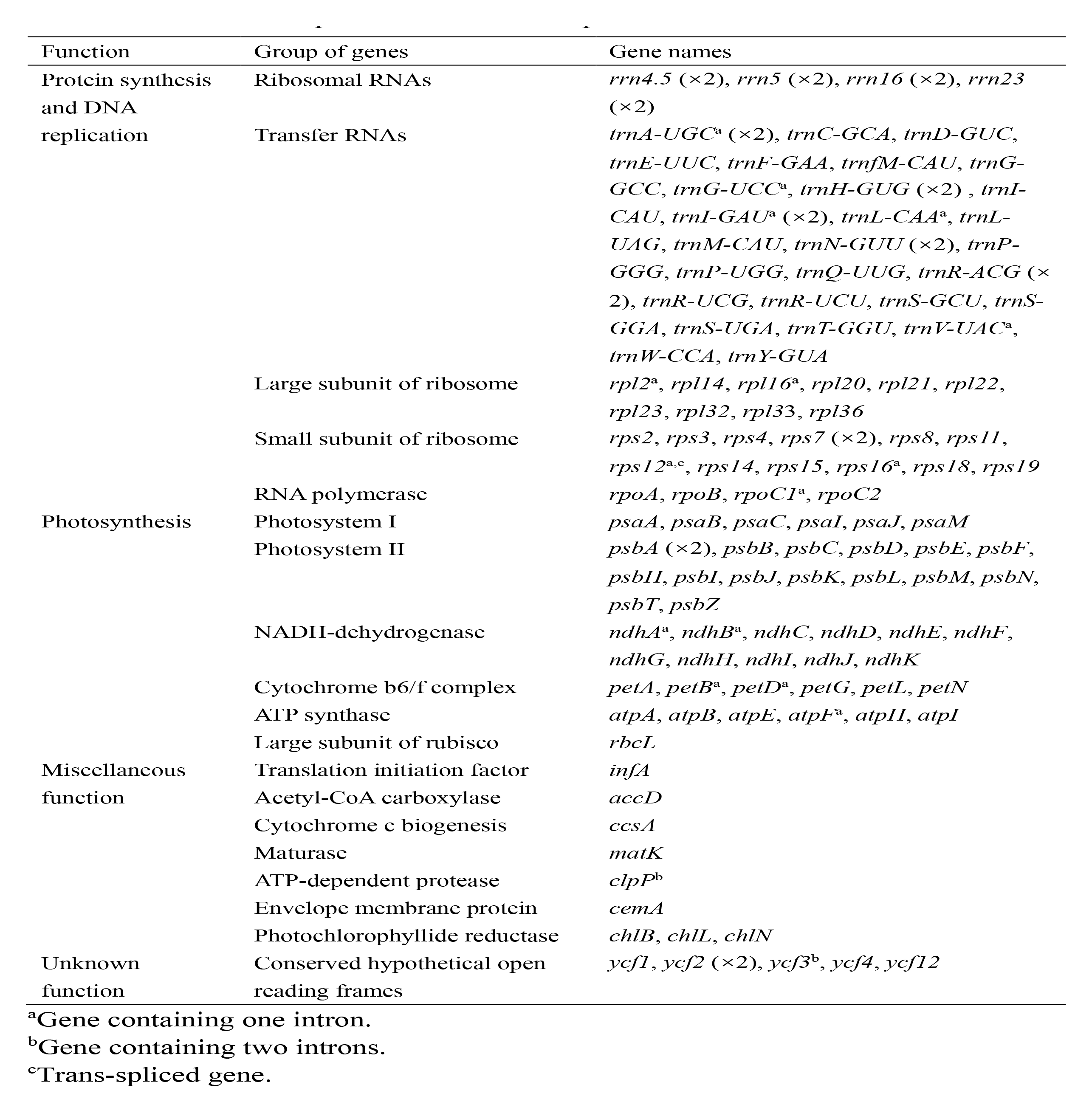
Genes in the plastome of *Cibotium* plants from China.

### Phylogenomic relationship within *Cibotium*

Alignments of whole length plastome and protein-coding genes were 136,298 bp and 73,080 bp, respectively. All three phylogenetic trees (Figures 5, S1) built with different datasets and methods showed generally similar topology and strongly supported the monophyly of *C. cumingii*, most *C. barometz*, as well as the five samples from NW Yunnan and NE Myanmar Yunnan-Myanmar. The five Yunnan-Myanmar samples formed one monophyletic clade (Clade A, Figure 5), which is sister to all the other accessions. The remaining samples of *C. barometz* except the one collected in Hainan island could be further divided into two lineages, i.e. Subclade E including samples from SE China (Zhejiang, Jiangxi, Fujian, Guangdong, Guizhou) and the Ryukyu Islands, and Subclade W including samples from SW China (Chongqing, Guangxi, Yunnan, Xizang). The Hainan sample clustered within Subclade E based on the protein-coding gene dataset with low support value (Figure 5), but formed a single clade sister to the combination of Subclade E and Subclade W (MLBS = 26) based on the whole length dataset (Figure S1). Therefore, both results failed to strongly resolve the topology among the southeastern and southwestern Subclades as well as the Hainan sample within *C. barometz*.

**FIGURE 5.**
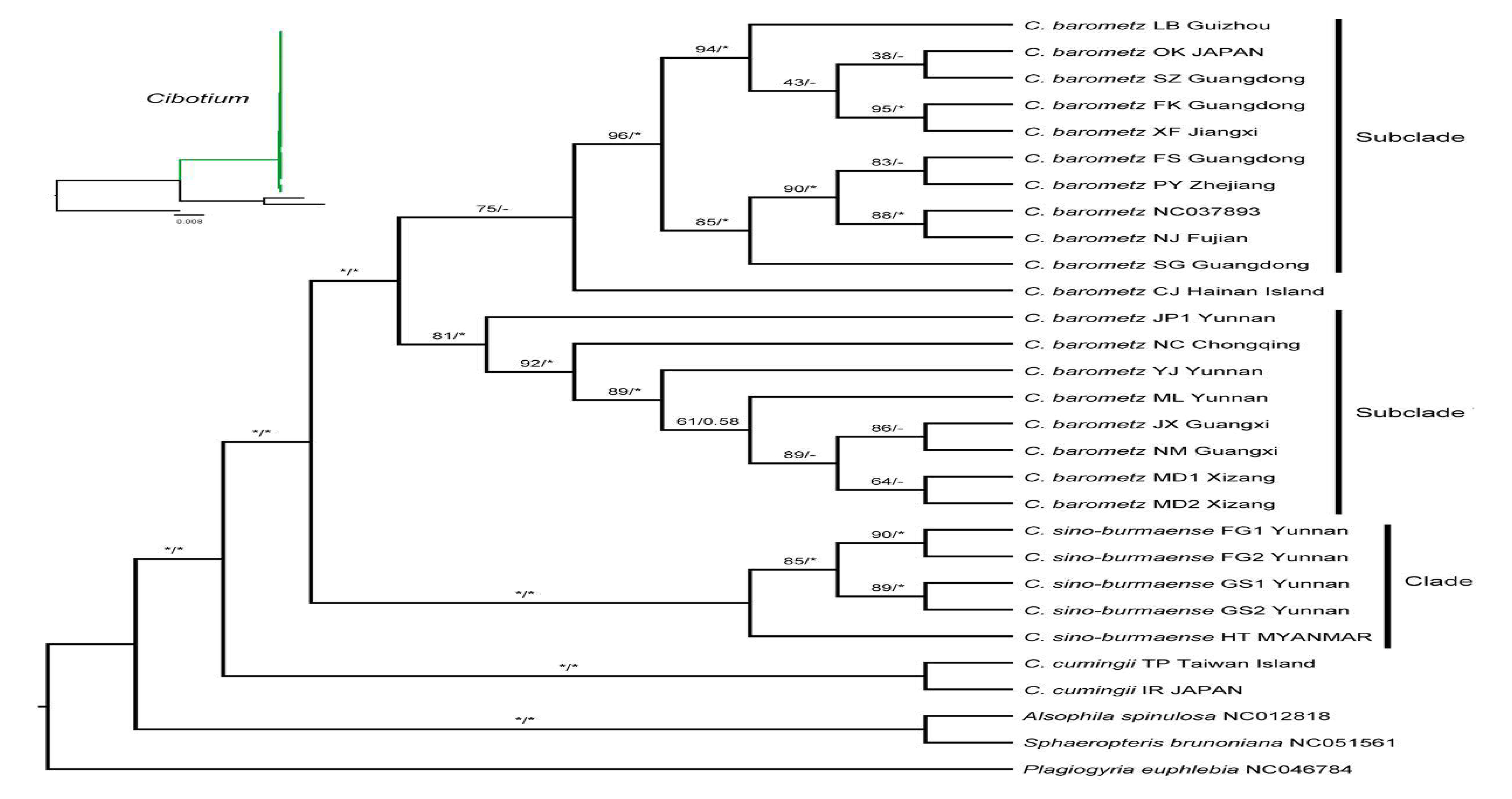
Maximum likelihood cladogram of the *Cibotium* plant from China and adjacent regions inferred from 85 concatenated protein-coding genes. Numbers above branches are bootstrap values (MLBS) and posterior probabilities (BIPP). Asterisk (*) indicates MLBS = 100% or BIPP = 1.0. En-dashe (–) indicates the lack of support value. The corresponding phylogram showing branch length is placed at the upper left corner.

### Genetic diversity and divergence of *Cibotium* species

Including two, five, nineteen, ten, and eight individuals, the estimated values of π in *C. cumingii*, *Cibotium* from Yunnan-Myanmar, *C. barometz*, as well as east and west lineages of *C. barometz* are 0.00032, 0.00008, 0.00027, 0.00020, and 0.00013 respectively. Among them, *C. cumingii* showed the richest genetic diversity though only two individuals were sampled, whereas the diversity is the lowest in the Yunnan-Myanmar *Cibotium* plants. Fixed difference numbers and Dxy values are as follows: *C. cumingii* & *C. barometz*, 183, 0.00175; *C. cumingii* & Yunnan-Myanmar *Cibotium*, 207, 0.00176; *C. barometz* & Yunnan

-Myanmar *Cibotium*, 80, 0.00085; Clade E & Clade W, 13, 0.00033. *C. cumingii* showed the greatest divergence level with two other species distributed in China. Interspecific variation of each species pair is significantly higher than lineages divergence within *C. borametz*.

### DNA Barcodes for *Cibotium* Species Discrimination

Based on phylogenetic topologies (Figure S2 A-I), only four (*trnL-trnF*, *trnG-trnR*, *rps4-trnS*, and *rbcL-accD*) among the nine traditional cpDNA loci are effective in the identification of *C. cumingii*, while merely the former two of the four could further discriminate Yunnan-Myanmar *Cibotium* correctly. None of them correctly showed the intraspecies divergence within *C. barometz*.

The nucleotide variability of *C. barometz* plastome was shown in Figure 6. Variable regions distributed evenly along plastome with π value less than 0.002 except a highly variable region within IR between *rrn16* and *rps12*. Five fragments (Figure 6) with moderate variation for species and lineage discrimination as well as suitable length and flanks for PCR amplification were chosen as candidate DNA barcode loci. Comparing with the nine old cpDNA loci, these new barcodes showed higher variability among the Chinese *Cibotium* species (Table 3), and were all capable to assign individuals of *C. cumingii*, Yunnan-Myanmar *Cibotium*, and two different lineages within *C. barometz* into respective clades correctly (Figure S2 J-N). The Hainan sample with uncertain phylogenetic position was clustered with samples of the western lineage by *rps3-rps19* and *<u>ndhA</u>*, but clustered into the eastern lineage by *chlB-trnQ*, *petD-rpoA* and *psaC-ndhG*.

**FIGURE 6.**
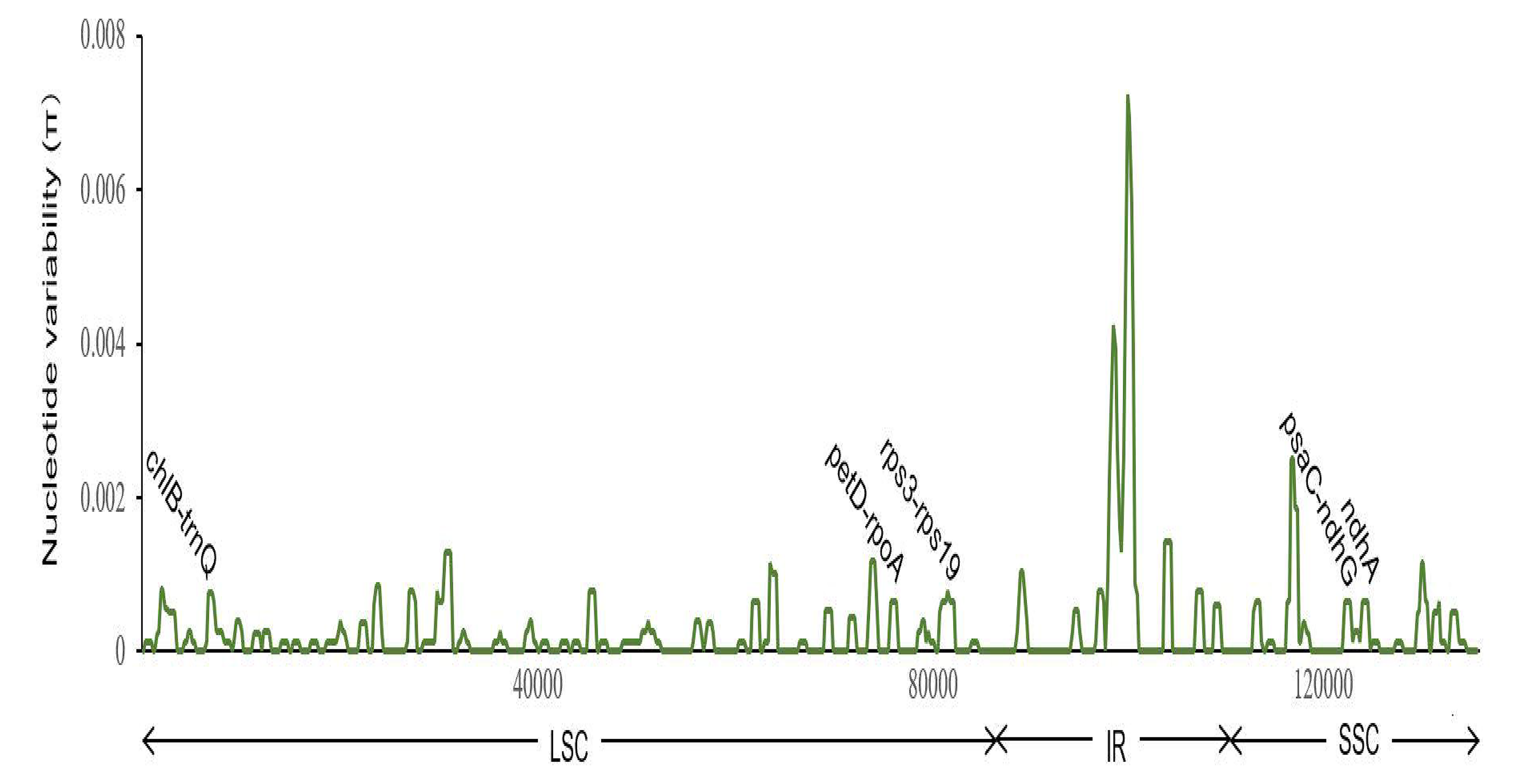
Sliding window analysis of 19 plastomes of *Cibotium barometz* (window length: 800 bp, step size: 200 bp). X and Y axes indicate the position of the midpoint of a window and nucleotide variability (π) of each window, respectively. Those marked fragments show the position of five newly designed DNA barcodes for inter-and intra-species discrimination in Chinese *Cibotium*.

**TABLE 3.**
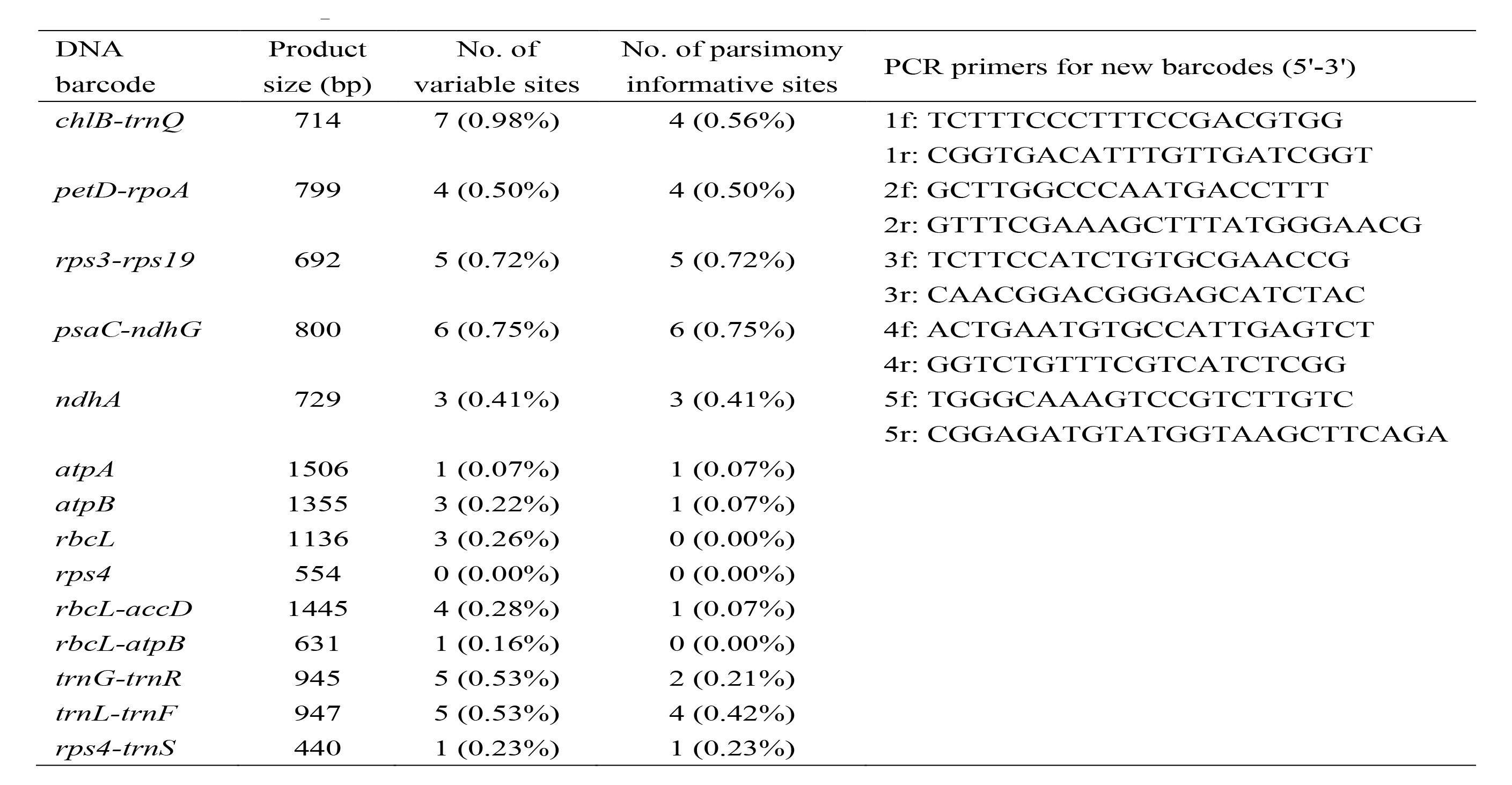
Characteristics of newly designed and traditional DNA barcodes of Chinese *Cibotium* plants.

## DISCUSSION

### *Cibotium* species from Yunnan-Myanmar

Based on our plastome-based phylogenetic relationship, *C. cumingii* is the species diverged firstly with all the other taxa of Chinese *Cibotium*. The result further distinguished two well-supported sister clades from the remaining samples, one compromising samples distributed in S China and the Ryukyu Islands corresponding to traditionally recognized *C. barometz*, the other compromising five samples from NW Yunnan and NE Myanmar (Clade A, Figure 5). Plastome characteristics including IR size and GC content as well as Dxy value also support the genetic difference of the Yunnan-Myanmar *Cibotium* from the widespread *C. barometz*. Therefore, we named these Yunnan-Myanmar samples as a new species, *Cibotium sino-burmaense*, hereafter.

We compared specimens of *C. sino-burmaense* with *C. barometz*, and found obvious differences of pinnules and sori characters (see details in the taxonomic treatment part). We checked spores of *C. sino-burmaense*, and found them shared similar perine features with other Asian species photographed by Gastony (1982) with strongly developed equatorial and distal ridges. However, the equatorial diameter of exospores is significantly larger (41-55 μm) than those of *C. barometz* from S China (30-45 μm). Additionally, we estimated the nuclear DNA content of our samples by flow cytometry with *Capsicum annuum* var. *annuum* (3.38 pg/C, Moscone et al., 2003) as the internal standard. Our results showed no incongruence with the record of *C. barometz* (4.58 pg/C, Clark et al., 2016) and detected no significant ploidy variation signal of *C. sino-burmaense* comparing with *C. barometz* (Figure S3). Our study also suggested two traditionally used cpDNA fragments (*trnL-trnF* and *trnG-trnR*) as well as five new barcodes that are highly effective in discriminating the new *Cibotium* species from other species of China. These genetic tools will be of benefit to the conservation of the Yunnan-Myanmar *C. sino-burmaense*, a cryptic species endemic to this region.

### Genetic divergence of *C. barometz* in China

In recent decades, integrating principles and methodologies such as taxonomy, phylogeny, and evolutionary ecology, has become a trend in aiding medicinal discovery, identification, and conservation (Sun et al., 2021; Xu et al., 2021; Zaman et al., 2021). Here, by means of phylogenomic analyses, we clarified the species boundary and presented the lineage divergence and geographic pattern of *C. barometz* in China, the highly demanded original resource of “Gouji”. Comparing with previous studies constrained in limited sampling areas (e.g., Wu et al., 2007; You and Deng, 2012), results of this study revealed the east-west divergence throughout the whole distribution region in S China. The geographic boundary is close to two general phylogeographic breaks of the Sino-Japanese floristic region, i.e. ca. 105°E and the boundary between the Second and Third ladders of landform in China as reviewed by Ye et al. (2017). Climate of the east and west sides of 105°E are dominated by Pacific and Indian monsoons respectively (Qiu et al., 2011), while altitude is significantly varied between and within different ladders (Li et al., 2013), which also shaped diverse ecological conditions (Fang et al., 2004). Heterogeneous climate and landform as well as refugia isolation resulted from intensity changes of monsoons may contributed to the east-west genetic split of *C. barometz* as demonstrated in other plant lineages (e.g., Bai et al., 2014; Sun et al., 2014; Kou et al., 2016). Additionally, the genetic difference also suggested that the east and west lineages should be concerned as at least two management units with respective genetic characters for conservation (Palsbøll et al., 2007). In the further, studies with population-level sampling of *C. barometz* and biparentally inherited nuclear genome data would evaluate within-population diversity on a finer scale, trace demographic history backwards, and predict the vulnerability of different lineages under the influence of habitat fragmentation and changing climate.

A large number of case studies have emphasized the fundamental role of germplasm resources played in high-quality genuine medicine production (e.g., Yao et al., 2020; Cheng et al., 2021; Xu et al., 2023). At present, all the cibotii rhizome slices sold on market come from natural sources without domestication, which varied seriously on medicinal quality (Ju et al., 2012; Yang et al., 2015). Environmental conditions affect the synthesis and accumulation of secondary metabolites which are usually medicinal components in plants (Li et al., 2020). Therefore, it is expected that *C. barometz* populations grow in habitats with diverged climate and ecology will also show pharmacodynamic difference. The diverged genetic background showed in this study will be beneficial to select specific high-quality germplasm resources from natural populations for cultivation, and elucidate the influence of multiple external factors on synthesis pathways of metabolites.

### Key to three *Cibotium* species of China

1a. Pinnules on basiscopic side of lower pinnae present or only one absent, rarely with two absent; sori 1–10 per pinnule segments 2
2a. Pinnules on acroscopic and basisicopic sides of a pinna nearly equal in length; apex of pinnule segments apiculate; sori oblong, usually 1–5 pairs per pinnule segment; average exospore equatorial diameter less than 43 μm 1. *C. barometz*
2b. Pinnules on basiscopic side of a pinna much shorter (c. 1/2) than those on the acroscopic side; apex of pinnule segments acute; sori oblong to spherical, usually 4–8 and sometimes over 10 pairs per pinnule segment; average exospore equatorial diameter more than 45 μm 2. *C. sino-burmaense*
1b. Pinnules on basiscopic side of lower pinnae usually three lacking; sori usually one or two per pinnule segment 3. *C. cumingii*

### Taxonomic treatment

(1) *Cibotium barometz* (L.) J. Sm., London J. Bot. 1 (1842) 437.
≡ *Polypodium barometz* L., Sp. Pl. 2 (1753) 1092.
≡ *Aspidium barometz* (L.) Willd., Sp. Pl., ed. 4 [Willdenow] 5 (1810) 268.
≡ *Nephrodium barometz* (L.) Sweet, Hort. Brit. [Sweet], ed. 2. (1830) 580.
≡ *Dicksonia barometz* (L.) Link, Fil. Spec. (1841) 166.
Type: —Not designated.
*= Cibotium assamicum* Hook., Sp. Fil. [W. J. Hooker] 1 (1844) 83, t.29B.
Holotype: —India. Assam, *Mrs. Mack s.n.* (in Sp. Fil. [W. J. Hooker] 1 (1844) t.29B).
= *Balantium glaucescens* Link, Fil. Spec. (1841) 40.
Type: —Not designated.
= *Cibotium glaucescens* Kunze, Farnkräuter 1 (1841) 63, t.31.
Holotype: —*s.coll. s.n.* (In Farnkräuter 1 (1841) 63, t.31.).
= *Dicksonia assamicum* Griff., Notul. 2 (1849) 607.

Lectotype (designated here): —India. Assam, *Griffith s.n.* (K barcode K001090393 [image!]).

Notes: —None original material of both basionyms, i.e., *Polypodium barometz* and *Balantium glaucescens*, was traced (see discussion by Holttum in *Fl. Malesiana*, ser. II, 1 (1963) 166).

Distribution: —China (Chongqing, Fujian, Guangdong, Guangxi, Guizhou, Hainan, Hunan, Jiangxi, Sichuan, Taiwan, Xizang, Yunnan, Zhejiang), Japan (Ryukyu Islands), Indonesia (Java to Sumatra), Malaysia, Myanmar, Thailand, Vietnam.

(2) ***Cibotium sino-burmaense*** X.C.Zhang & S.Q.Liang, sp. nov. (Figure 7)

**FIGURE 7.**
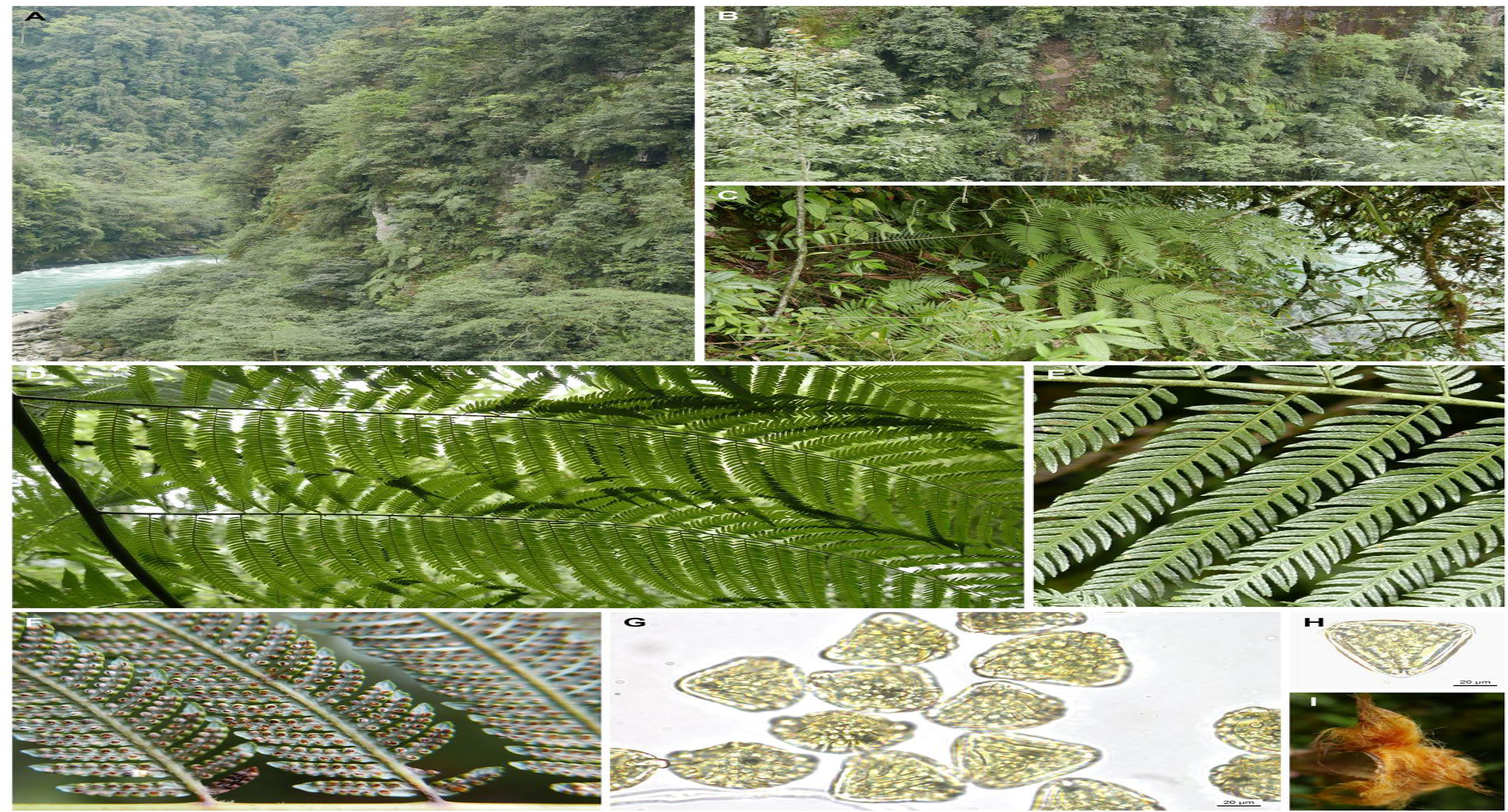
Habitat and morphology of *Cibotium sino-burmaense* sp. nov. from Dulongjiang, Gongshan, Yunnan, China. **(A-B)** Habitat. (**C**) Habit. (**D**) Pinnae on abaxial side. (**E**) Pinnules on adaxial surface. (**F**) Pinnules with opened sori. (**G**-**H**) Spores under light microscopy. **(I)** Golden filiform hairs on stipe base. Photographs by, X.-C. Zhang (A–F & I), and S.-Q. Liang (G–H).

Diagnosis: —This new species resembles *C. barometz* and *C. cumingii*, differing from the former in the significantly shortened pinnule length on basiscopic side, as well as acute apex and more sori of pinnule segments, and from the latter in the denser sori per pinnule segment and presence of the second and third pinnules on the basiscopic side of lower pinnae.

Holotype: —**China.** Yunnan: Gongshan county, Dulongjiang Township, 2 May 2022, *X.C. Zhang 12880* (PE).

Note: —The holotype consists of a single large frond mounted on fifteen herbarium sheets, labelled “sheet 1” to “sheet 15”.

Description: —Rhizome prostrate, stout, densely covered with shiny yellowish brown long hairs. Stipes thick, up to 80 cm or more, dark brown to purplish black at base and becoming green upwards, covered with long hairs similar to those on rhizome at base, upper part covered with small, appressed flaccid hairs. Lamina ovate, 2-pinnate-pinnatifid, up to 3 m, subleathery, adaxial surface deep green, abaxial surface glaucous, with small flaccid hairs on midrib; pinna 8–10 pairs, alternate, stalked, medial pinnae 60–80 × 20–30 cm, basal pinna pairs reduced slightly; pinnules more than 30 pairs per lower pinna, shortly stalked, up to 20 cm on the acroscopic side, 10-14 cm on the basiscopic side; pinnule segments, alternate, slightly falcate, with acute apex, margins crenulate to serrulate-serrate. Sori oblong to spherical, usually 4–8 and sometime over 10 pairs at base of lower pairs of pinnule segments; indusia bivalvate, outer indusia larger, orbicular, inner significantly smaller, oblong. Spores pale yellowish, with strongly developed equatorial and distal ridges.

Etymology: —*Sino-burmaense* is derived from the known distribution of this species along China-Myanmar border.

Additional Specimens Examined: —**China.** Yunnan: Gongshan county, Dulongjiang Township, 23 Jan 2017, *X.C. Zhang & al. 8134*; Fugong County, 26 Apr 2022, *X.C. Zhang 12831.* **Myanmar.** Kachin: Htawgaw, Apr 1925, *G. Forrest 26496* (PE barcode 01654827, 01654828, 00388348).

Distribution and habitat: —China (NW Yunnan), Myanmar (Kachin). On cliff with open canopy.

(3) *Cibotium cumingii* Kunze, Farrnkräuter 1 (1841) 64, 65.

≡ *Cibotium barometz var. cumingii* (Kunze) C. Chr., Index Filic. 3 (1905) 183.

Lectotype (designated here): —Philippines. Luzon, *H. Cuming 123* (K barcode K000376224 [image!]; isolectotypes: K barcode K000376225 [image!], K000376228, K000376229, K000376231, K000376232; BM barcode BM001048122 [image!]; E barcode E00822366 [image!], E00822367 [image!], E00822369 [image!], E00822373 [image!]; P barcode P00633260 [image!], P00633261 [image!], P00633262 [image!]; US barcode 00134826 [image!]; Z barcode Z-000002072 [image!]).

= *Cibotium crassinerve* Rosenst., Meded. Rijks-Herb. 31 (1917) 4.

Lectotype (designated here): —Philippines. Luzon, Benguet, Dec 1908, *H. M. Curran & M. L. Merritt 15800* (L barcode L 0051165 [image!]; isolectotype: MICH No. 1190172 [image!]).

= *Cibotium taiwanense* C.M.Kuo, Taiwania 30 (1985) 56, 57.

Lectotype (designated here): —China. Taiwan, Hsinchu, Chu-tong, Aug 1972, *C. M. Kuo 1703* (TAI No. 149443 [image!]; isolectotypes: TAI No. 148725 [image!], 150173 [image!]).

Distribution: —China (Taiwan), Japan (Ryukyu Islands), Phillipines.

## CONCLUSION

This study presented conserved structure and gene composition of chloroplast genome within *Cibotium* from China. Based on phylogenomic analyses, we constructed a well-supported phylogeny of Chinese *Cibotium*, and indicated that there are three species distribute in China, namely *C. barometz*, *C. cumingii*, and *C. sino-burmaense*, an overlooked cryptic species from the NW Yunnan and NE Myanmar. Moreover, our results uncovered the east-west lineage divergence in *C. barometz*. We also evaluated the species resolution of nine old cpDNA loci, and suggested five new cpDNA barcodes which are capable to identify all the above-mentioned species and lineages of Chinese *Cibotium* accurately. In conclusion, our findings will improve people’s understanding on the germplasm resource diversity of this endangered medicinal plant group, and play a guiding role in its wild population conservation and medical value exploitation.

## DATA AVAILABILITY STATEMENT

## AUTHOR CONTRIBUTIONS

X-CZ, K-XL and R-HJ designed this study. FW, L-MT, BQ, Y-YC and Y-HH collected and cultivated plant materials of this study. S-QL performed experiments, analyzed the data, and wrote the manuscript.

## FUNDING

This research was supported by “Evaluation of the Germplasm Resources of the Protected Plant *Cibotium barometz*” project (GuiLinYan[RC]2302) of Guangxi Forestry Research Institute, the National Plant Specimen Resource Center Project (NPSRC) (E0117G1001), “Field Survey and Conservation Studies of some State Key Protected Fern Species” project in National Forestry and Grassland Administration, “2022 Central Financial Forestry Reform and Development Funds: Collection, Conservation and Use of Germplasm Resources of *Cibotium barometz*” of Department of Forestry of Guangxi Zhuang Autonmous Region, as well as “Survey and Collection of Germplasm Resources of Woody and Herbaceous Plants in Guangxi, China” (GXFS-2021-34).

## ACKNOWLEDGMENTS

We appreciate Dr. Xiang-Yun Zhu for valuable discussion about taxonomic treatment. We thank Dr. Jie Yang, Mr. Jun-Yong Tang and Mr. Ji-Gao Yu for their help in plastome data analyses. We also thank Jin-Dan Zhang from the Plant Science Facility of the Institute of Botany, Chinese Academy of Sciences for the technical assistance on flow cytometry.

## Notes

### Competing Interest Statement

The authors have declared no competing interest.

