## Supplemental figures for "Plastome-based Phylogenomic analyses provide insights into the germplasm resource diversity of *Cibotium* in China"

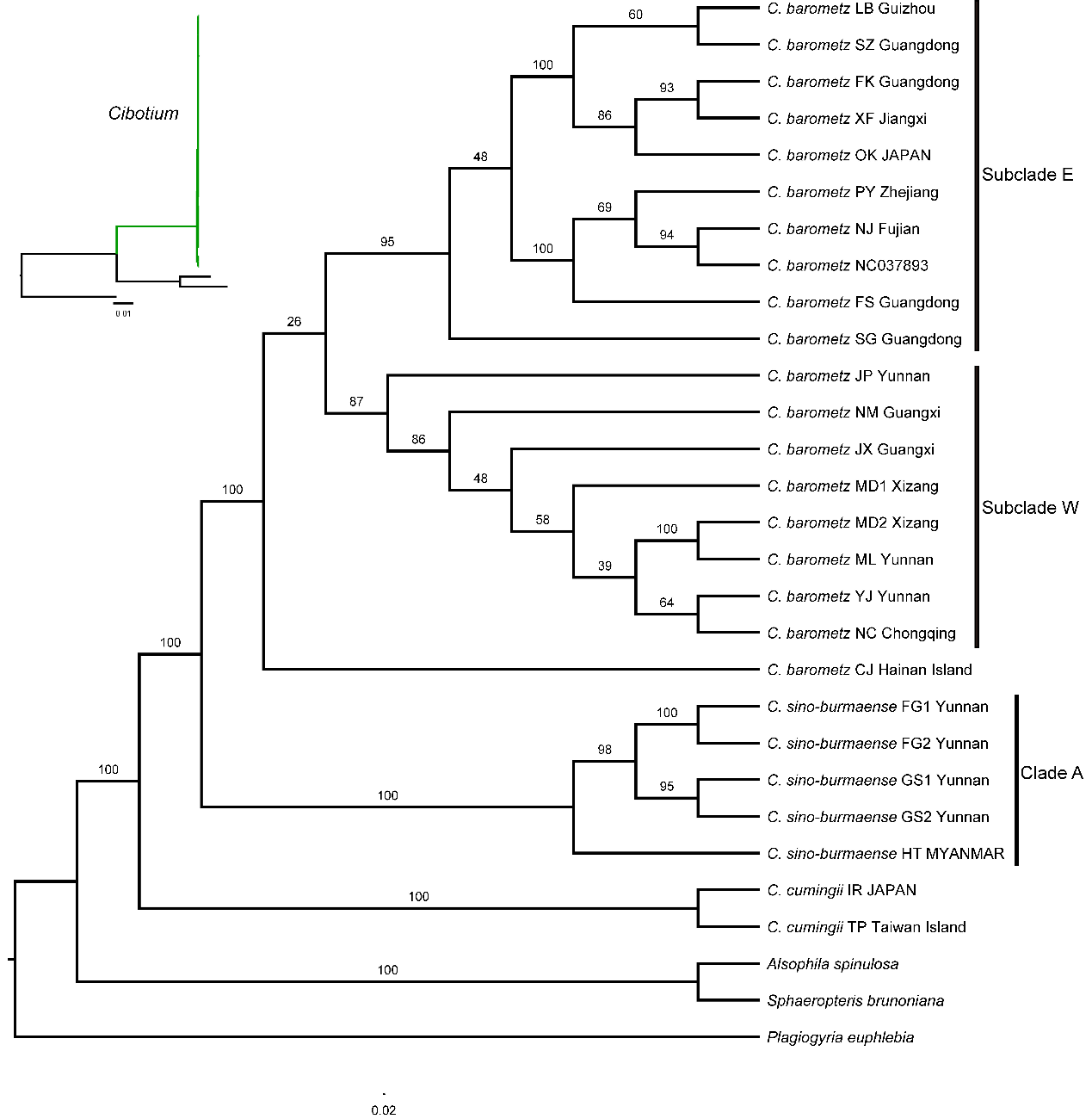

**FIGURE S1** Maximum likelihood cladogram of the *Cibotium* plant from China and adjacent regions inferred from whole length plastomes. Numbers above branches are bootstrap values (MLBS). The corresponding phylogram showing branch length is placed at the upper left corner.

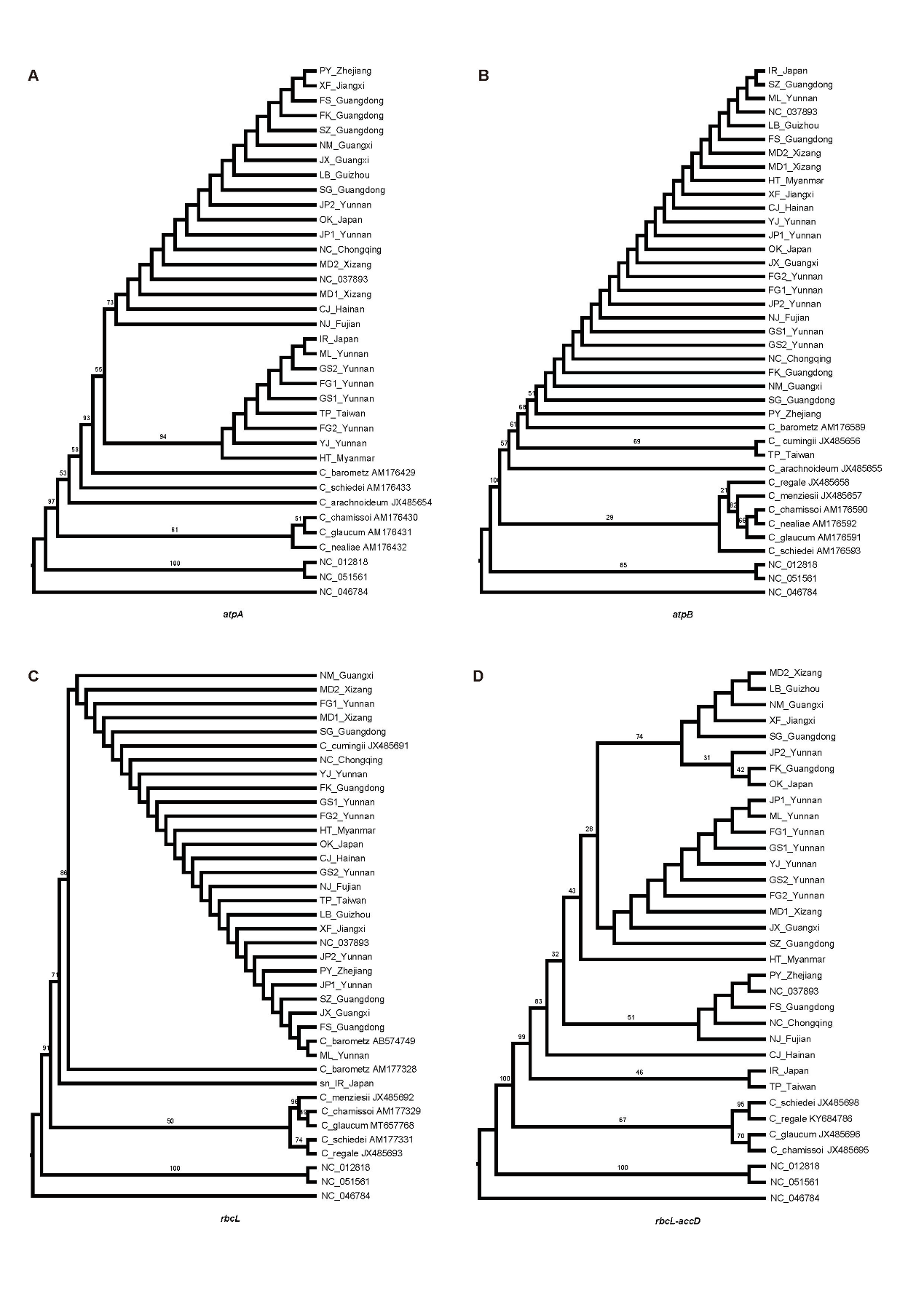

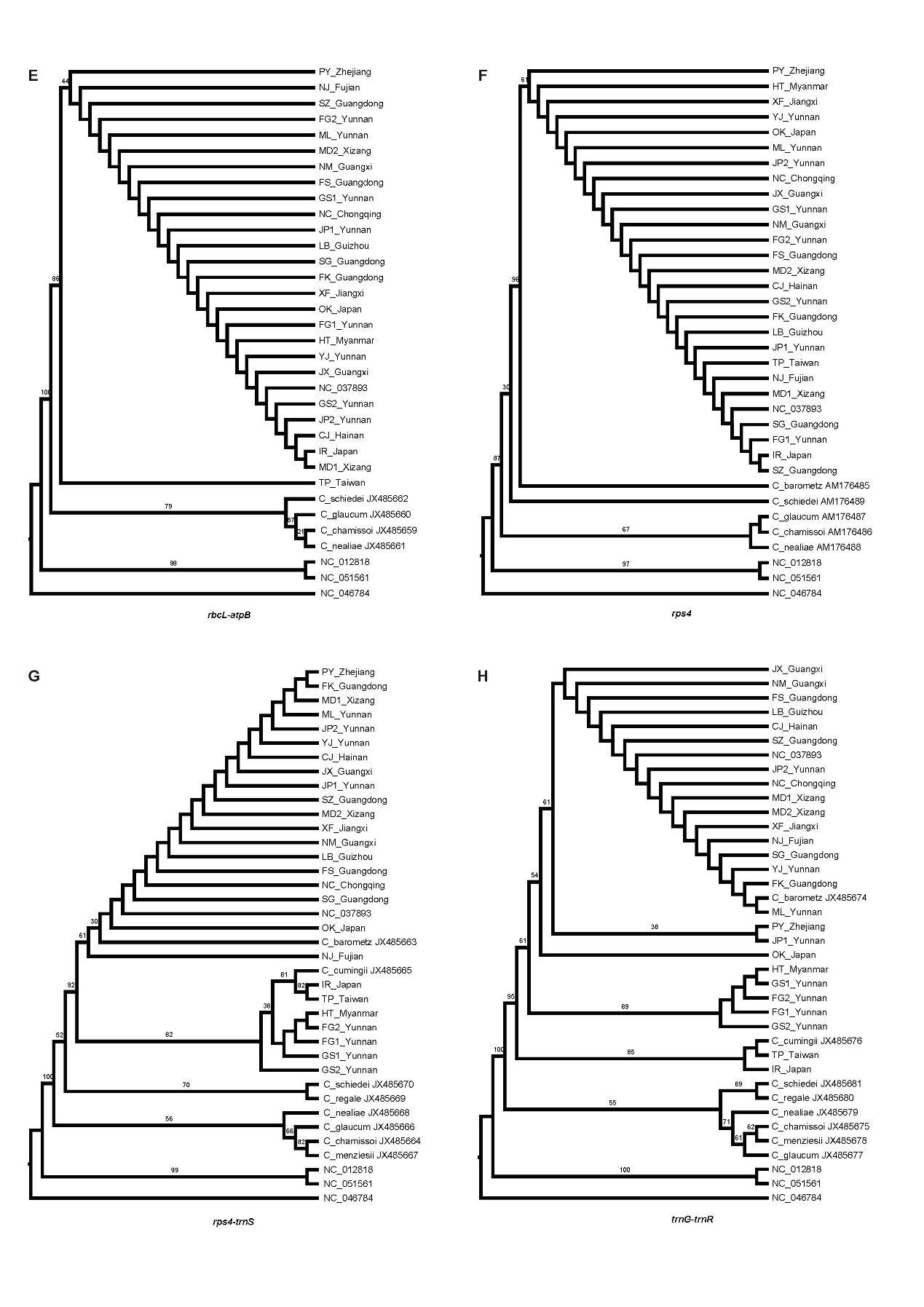

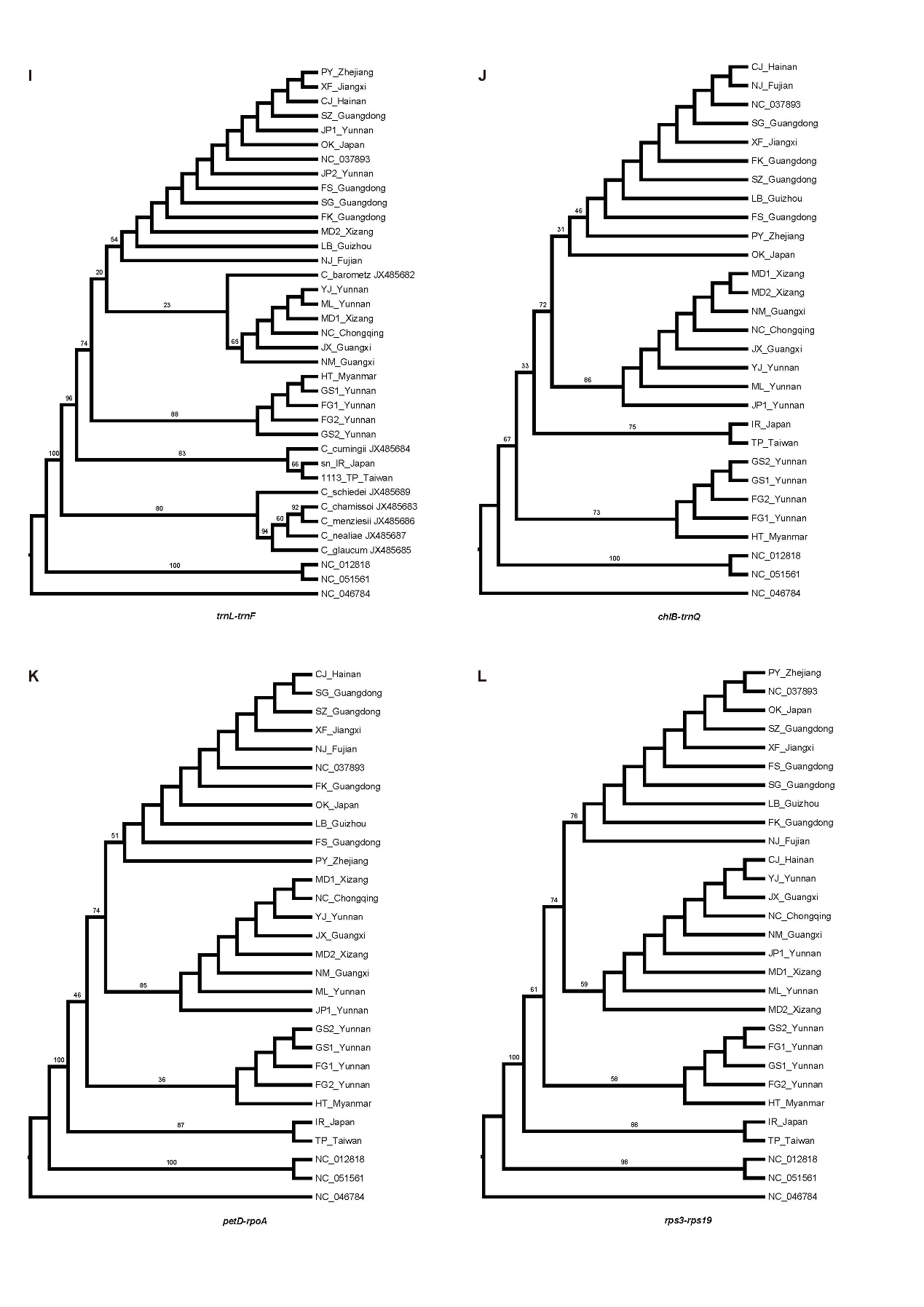

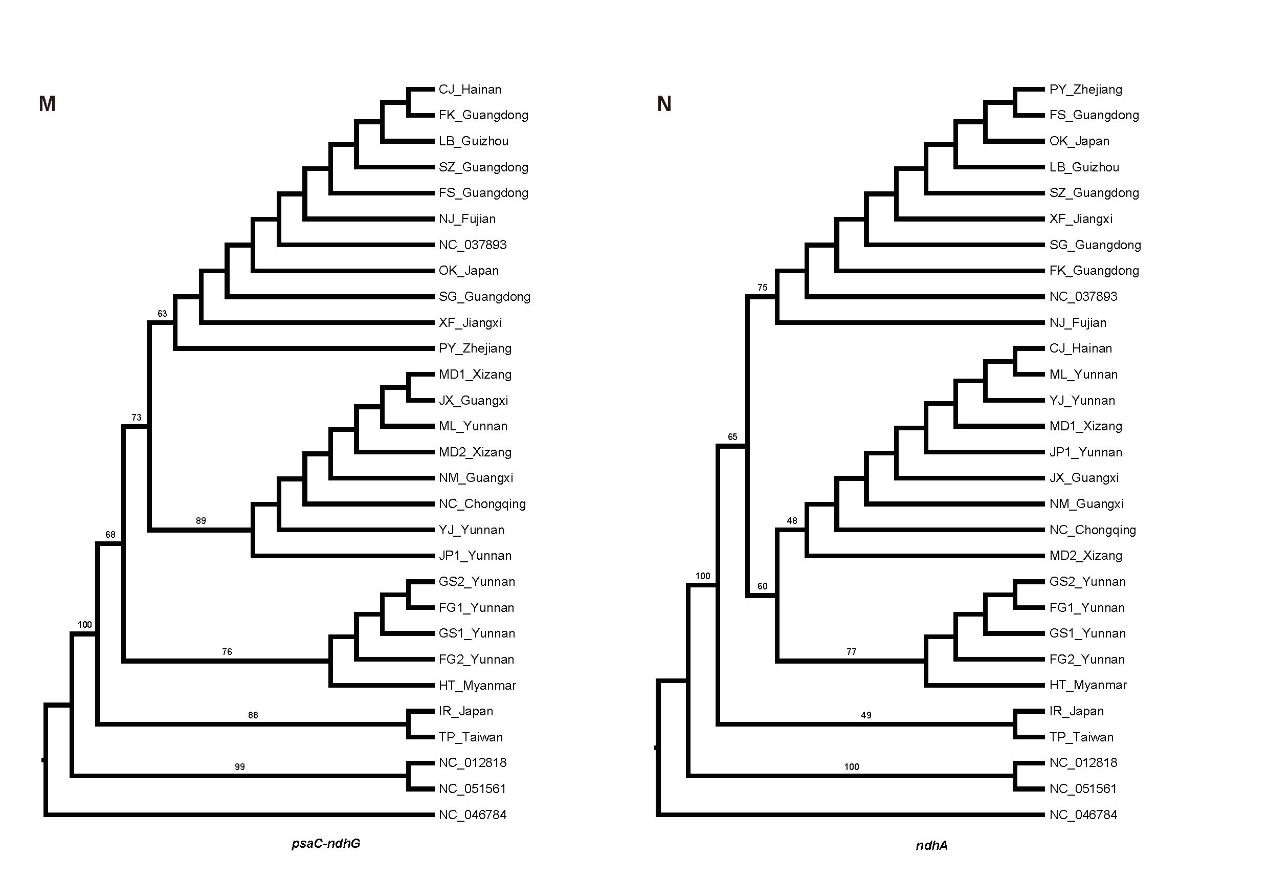

**FIGURE S2** Maximum likelihood cladogram of the *Cibotium* built with different cpDNA barcode loci. Numbers above branches are bootstrap values.

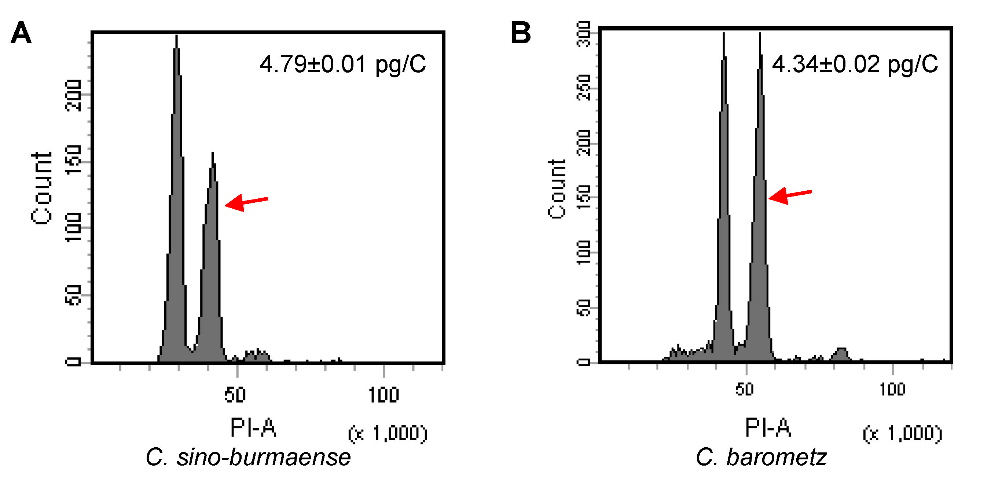

**FIGURE S3** Histograms of fluorescence intensity in two *Cibotium* species. The peak marked with a red arrow represents *Cibotium* sample, while the other peak represents the internal standard *Capsicum annuum* var. *annuum* (nuclear DNA content: 3.38 pg/C).
